# mTORC2 Interactome and Localization Determine Aggressiveness of High-Grade Glioma Cells through Association with Gelsolin

**DOI:** 10.1101/2022.10.12.511677

**Authors:** Naphat Chantaravisoot, Piriya Wongkongkathep, Nuttiya Kalpongnukul, Narawit Pacharakullanon, Pornchai Kaewsapsak, Chaiyaboot Ariyachet, Joseph A. Loo, Fuyuhiko Tamanoi, Trairak Pisitkun

**Author notes:** Equal first authors (Naphat Chantaravisoot, Piriya Wongkongkathep). Corresponding authors: Naphat Chantaravisoot and Trairak Pisitkun Send correspondence to: Naphat Chantaravisoot, Dept. of Biochemistry, Faculty of Medicine, Chulalongkorn University, 1873 Rama IV Pathumwan Bangkok, 10330, Thailand.

## Abstract

The mTOR complex 2 (mTORC2) has been implicated as a key regulator of glioblastoma cell migration. However, the roles of mTORC2 in the migrational control process have not been entirely elucidated. Here we elaborate that active mTORC2 is crucial for GBM cell motility. Inhibition of mTORC2 impaired cell movement and negatively affected microfilaments and microtubules’ functions. We also aimed to characterize important players involved in the regulation of cell migration and other mTORC2-mediated cellular processes in GBM cells through the proteomic and bioinformatic analyses. Therefore, we quantitatively characterized the alteration of mTORC2 interactome under selective conditions using affinity-purification mass spectrometry (AP-MS) in glioblastoma. We demonstrated that changes in cell migration ability specifically altered mTORC2-associated proteins. Gelsolin (GSN) was identified as one of the most dynamic proteins. The mTORC2-GSN linkage was mostly highlighted in high-grade glioma cells and shown to connect functional mTORC2 to multiple proteins responsible for directional cell movement in GBM. GSN also mediated membrane localization of mTORC2. Loss of GSN disconnected mTORC2 from numerous cytoskeletal proteins but did not affect mTORC2 integrity. In addition, we reported 86 stable mTORC2 interactome involved with diverse molecular functions, predominantly cytoskeletal remodeling in GBM. Our findings help expand future opportunities for predicting the highly migratory phenotype of brain cancers in the clinical investigation.

## INTRODUCTION

According to the 2016 World Health Organization (WHO) classification system, glioblastoma multiforme (GBM) has been classified as a grade IV type of diffuse astrocytic tumors and divided into subgroups depending on their molecular profiles (Louis et al., 2016). GBM is the most malignant type of gliomas and one of the most deleterious human cancers based on low overall survival (OS) of around 15 months, estimated 10-year survival rate of 0.71%, high recurrence rates after diagnosis, and resistance to both chemotherapy and radiotherapy. Despite multiple aggressive treatments, most cases experience recurring tumors and finally progress to death. (DeAngelis, 2001; Masui et al., 2016; Tykocki and Eltayeb, 2018).

The mechanistic target of rapamycin complex 2 (mTORC2) has been implicated as one of the major signaling molecules in various types of brain cells and an attractive therapeutic target for GBM (Masui et al., 2013; Thomanetz et al., 2013; Wu et al., 2014). The mTORC2 mainly consists of mTOR kinase, RICTOR, MAPKAP1 (or mSIN1), and MLST8 (Laplante and Sabatini, 2012). Growth factors can stimulate mTORC2 resulting in AKT phosphorylation at serine 473 (Sarbassov et al., 2004). This multiprotein complex has been identified as a crucial regulator of actin cytoskeleton reorganization through PKCα phosphorylation and other PKC isoforms (Fu and Hall, 2020; Jacinto et al., 2004; Oh and Jacinto, 2011).

The relationship between deregulated mTORC2 signaling pathway and the malignancy of gliomas has been increasingly reported as more mTORC2 functions are revealed (Das et al., 2011; Masui et al., 2014; Masui et al., 2017). Hyperactivation of mTORC2 could also be observed in clinical samples from multiple cancers in various studies (Bian et al., 2015; Guertin et al., 2009; Jiang et al., 2017; Magee et al., 2012; Sticz et al., 2017; Tanaka et al., 2011). Moreover, the inhibition of mTORC2 by several ATP-competitive mTOR kinase inhibitors and RICTOR/mTORC2 depletion could decrease cell growth, proliferation, motility, invasiveness, and stemness properties of GBM cells and malignant glioma tumors (Chantaravisoot et al., 2015; Gini et al., 2013; Li et al., 2020; Mecca et al., 2018). Dysregulated mTORC2 signaling has been associated with the metabolic reprogramming events in GBM (Gu et al., 2017; Masui et al., 2015). The mTORC2 could phosphorylate FLNA at its stabilizing residue (S2152), promoting migration and invasion of GBM cells through the interactions of FLNA and integrins at the plasma membrane (Chantaravisoot *et al*., 2015; Sato et al., 2016). Nevertheless, the mechanisms of how mTORC2 may interact with other proteins to directly control cancer cell motility have not been thoroughly investigated.

Proteomic studies of GBM have been extensively performed in several aspects, mainly to determine the biomarkers (Pirlog et al., 2019; Silantyev et al., 2019). Quantitative proteomics has become a powerful method for identifying valid predictive biomarkers for brain cancers (Miyauchi et al., 2018). Moreover, mass spectrometry has emerged to characterize protein-protein interactions (PPIs) in disease or drug perturbations (Smits and Vermeulen, 2016). However, none of the proteomic studies in GBM focuses explicitly on its highly migratory characteristics or mTORC2-mediated directional migration of the cancer cells.

Here, we aim to extensively decipher the mTORC2 interactome and the mechanisms underlying how mTORC2 promotes highly migratory characteristics of GBM cells. Using quantitative proteomics and systems biology approaches, we demonstrate that mTORC2 interacts with all types of the cytoskeleton and its cellular localization indicates cancer cells’ ability to promote migration. Comparative analyses of the proteomic data sets of high- and low-grade glioma cells were combined to screen for vital regulators responsible for glioblastoma’s enhanced motility. Finally, we identify GSN as one of the distinct players connecting mTORC2 to several actin-binding and microtubule-associated proteins. Our work comprehensively provides proven evidence of direct and indirect PPIs between mTORC2 and the cytoskeleton-associated proteins that could generate the complex network initiating the regulation of cytoskeletal dynamics and help expand future opportunities for predicting the highly migratory phenotype of brain cancers in the clinical investigation.

## RESULTS

### mTORC2 is a **Crucial R**egulator of **G**lioblastoma **M**igration

To confirm the regulation of mTORC2 signaling cascade in GBM cells, we performed the western blotting analysis of the U87MG whole-cell lysate to show that growth factors can substantially activate the mTORC2 signaling in cells by inducing the pAKT (S473) while amino acids did not elevate pAKT level. In contrast, both serum and amino acids could stimulate mTORC1 signaling by increasing the level of pS6 (S235/236). Also, we showed that AZD8055, an ATP-competitive mTOR inhibitor, could inhibit both mTORC1 and mTORC2 by significantly reducing pS6 and pAKT levels, respectively. Nevertheless, rapamycin, a conventional mTOR inhibitor, potently inhibited only mTORC1 (Figure 1A). Then, we investigated the dose-dependent effects of AZD8055 on the phosphorylation of mTORC2 downstream targets. We found that amounts of pFLNA (S2152), pAKT (S473), and pS6 (S235/236) were significantly decreased at concentrations above 0.2 µM, and the most effective condition was at 2.0 µM (Figure 1B). Since AZD8055 is highly specific to mTOR kinase at concentrations up to 10 µM (Chresta et al., 2010), we selected the treatment condition at 2.0 µM as the main concentration for all experiments. We showed that RICTOR siRNA treatment decreased the levels of activated mTORC2 effectors, pFLNA, and pAKT (Figure 1C). Next, to emphasize the significance of mTORC2 signaling in GBM cell migrational control, we observed the migration ability of U87MG cells under different culturing conditions. U87MG cells under serum starvation, treated with AZD8055 or siRICTOR, showed impaired migration than cells in the activated state, while rapamycin did not decrease the migration rate (Figures 1D and 1E). The results suggested that mTORC2 plays a primary role in supporting GBM cell motility.

**Figure 1.**
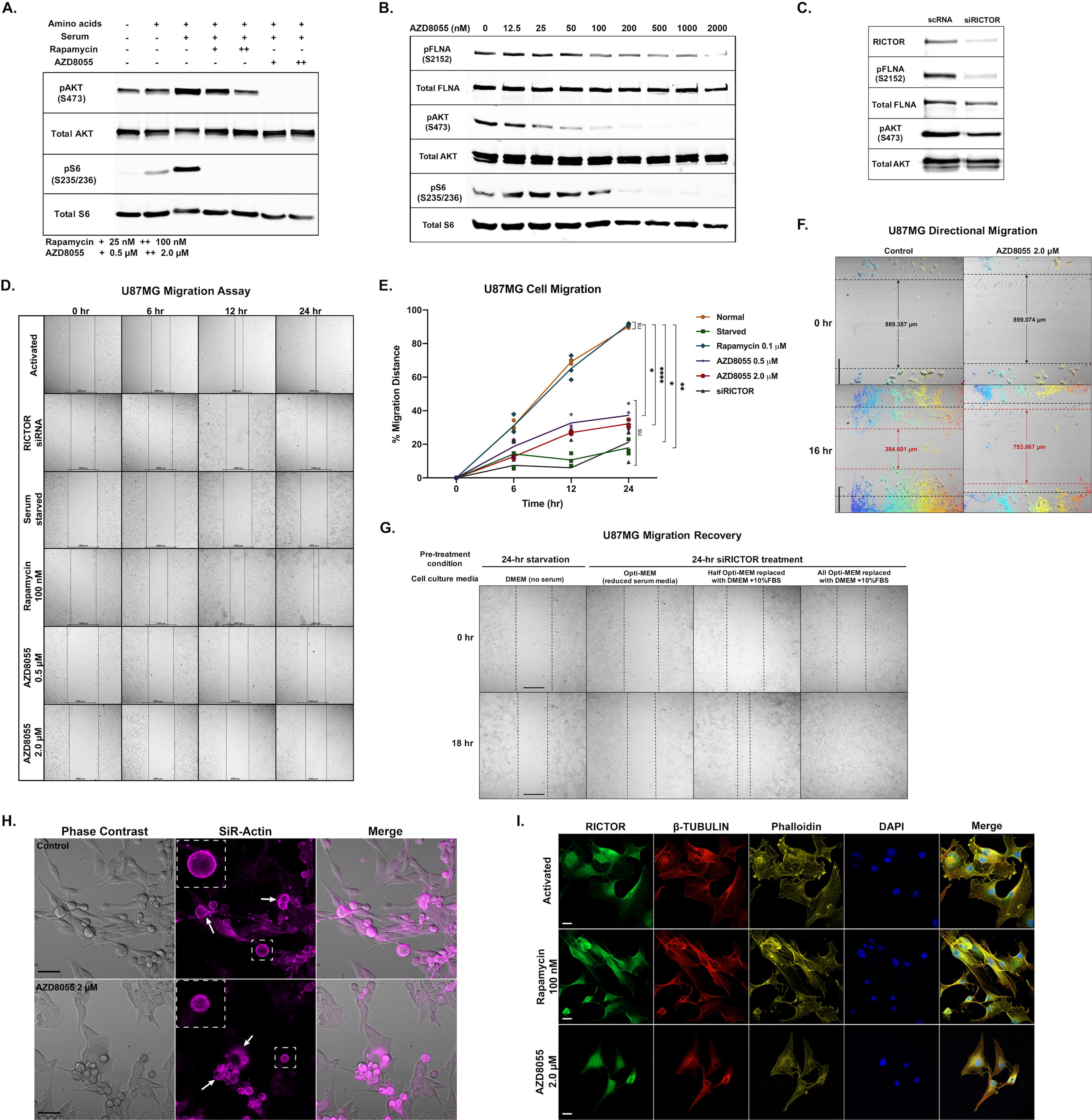
mTORC2 Signaling Pathway and Cell Migrational Control in U87MG. (A) Western blot showing activation of mTORC1 (pS6) and mTORC2 (pAKT) by amino acids and serum, and inhibition by rapamycin and AZD8055. (B) Western blot showing dose-dependent inhibition of mTORC1 (pS6) and mTORC2 (pFLNA and pAKT) by AZD8055. (C) Western blot analysis of RICTOR, pFLNA, and pAKT showing mTORC2 kinase activities after *RICTOR* knockdown (D) Wound-healing migration assay of U87MG cells under activation and inhibition (siRICTOR, serum-starved, 100 nM rapamycin, 0.5 µM and 2.0 µM AZD8055) conditions at 0, 6, 12, and 24 hr. Scale bar, 100 µm (E) Migration distance percentage quantification of U87MG cells under different culturing conditions shown in (D); n = 3. Statistical significance was calculated at the endpoint (24 hr) using two-way ANOVA with Tukey’s multiple-comparisons test comparing each treated group with the normal group: * p < 0.05, ** p < 0.01, *** p < 0.001, **** p < 0.0001, ns = not significant. (F) Directional migration (16 hr) of normal and 2.0 µM AZD8055-treated U87MG cells. Colored-tracking lines represent migrating direction of each cell in a circle. Scale bar, 200 µm. See also Video S1. (G) Cell migration recovery by serum supplementation (18 hr) of siRICTOR-treated cells. Scale bar, 100 µm. See also Video S2. (H) Snapshot from live-cell imaging of U87MG cells containing actin labeled with SiR-Actin under normal and AZD8055-treated conditions. See also Video S3. (I) Immunofluorescence staining of F-actin and the microtubules of activated, rapamycin-treated (100 nM, 24 hr), and AZD8055-treated (2.0 µM, 24 hr) U87MG cells. Scale bar, 20 µm.

Moreover, we performed live-cell imaging to investigate cell migration behavior when mTORC2 was blocked. The wound-healing assay results showed that cells treated with AZD8055 could not close the gap efficiently, and their directional migration was disrupted, as shown by the tangled cell tracking lines (Figure 1F; Video S1). Subsequently, 24-hr *RICTOR*-knocked down and serum-starved cells were tested to investigate whether activating mTORC2 complexes by growth factors could restore cells’ migration ability. We replaced the serum-deprived culture media with DMEM containing 10% FBS and found that the cancer cells closed the gap within 18 hours after media replacement. Conversely, serum-starved cells had not acquired the migration capability (Figure 1G; Video S2). Our findings have concluded that active mTORC2 is critical for the migration of GBM cells.

### Inactivation of mTORC2 Affects Actin Cytoskeleton and Microtubule Network Architecture and Dynamics

To better understand the causes of GBM cells’ impaired migration after the mTORC2 activity was disrupted, we first investigated the actin cytoskeleton dynamics when cells were treated with AZD8055 by live-cell imaging (Figure 1H; Video S3). The results showed that when inhibiting mTORC2 complexes, the actin network was highly affected, leading to abnormal migration. We observed significantly less actin filament surrounding the AZD8055-treated cell membrane region. Therefore, we hypothesized that the loss of organized actin networks near the plasma membrane (PM) disabled cells from moving regularly. In addition, we proposed that microtubule dynamics might also be defective when migrational control became strongly inhibited under mTORC2-suppressing conditions. We aimed to explore the effects on the overall structures of the actin cytoskeleton and the microtubules using live-cell fluorescence imaging. U87MG cells treated with AZD8055 and siRICTOR, except for rapamycin, showed a considerably non-functioning, abnormal cytoskeleton. Apart from damaged actin filaments, the shrinkage of microtubules was also observed. The fluorescently labeled β-tubulin proteins appeared mainly in the perinuclear area, suggesting that cells could not extend their microtubules to the PM. (Figure S1; Video S4 and S5). We suspected that active mTORC2 might interact with tubulin isoforms and microtubule-associated proteins to control microtubule dynamics.

We further confirmed the observation by staining F-actin and β-tubulin with RICTOR proteins. We discovered that RICTOR localization altered from a concentrated amount near the PM in the activated cells to nuclear and perinuclear regions in AZD8055-treated cells. In contrast, GBM cells treated with rapamycin did not exhibit any distinct changes (Figure 1I). Overall results indicated that active mTORC2 might promote cell migration by physically interacting with microtubules, microfilaments, and cytoskeletal accessory proteins.

### mTORC2 Complex Integrity is Critical for Its Localization

Since RICTOR colocalized with FLNA in GBM cells and mTORC2 inhibition caused FLNA dissociation from the actin cytoskeleton (Chantaravisoot *et al*., 2015), we hypothesized that the localization of RICTOR and FLNA would be changeable depending on the mTORC2’s activation status. We found that RICTOR and FLNA localized differently in starved and AZD8055-treated compared to the activated cells (Figure 2A). We later explored whether RICTOR could represent the whole mTORC2 complex. The immunofluorescence (IF) staining of the main components of mTORC2; mTOR, RICTOR, and MAPKAP1 exhibited that all proteins colocalized near the cell membrane under the mTORC2-activating state. In contrast, starvation and AZD8055-treated conditions resulted in fewer PM-linked mTORC2 complexes. Also, the three proteins only partially colocalized in both inactivation conditions (Figure 2B). However, rapamycin did not affect RICTOR-MAPKAP1 colocalization. Our results correlated with a previous study that rapamycin does not affect mTORC2 assembly (Rosner and Hengstschlager, 2008). Hence, the results suggested that the functionally active mTORC2, associated with a highly migratory phenotype, is localized at the PM.

**Figure 2.**
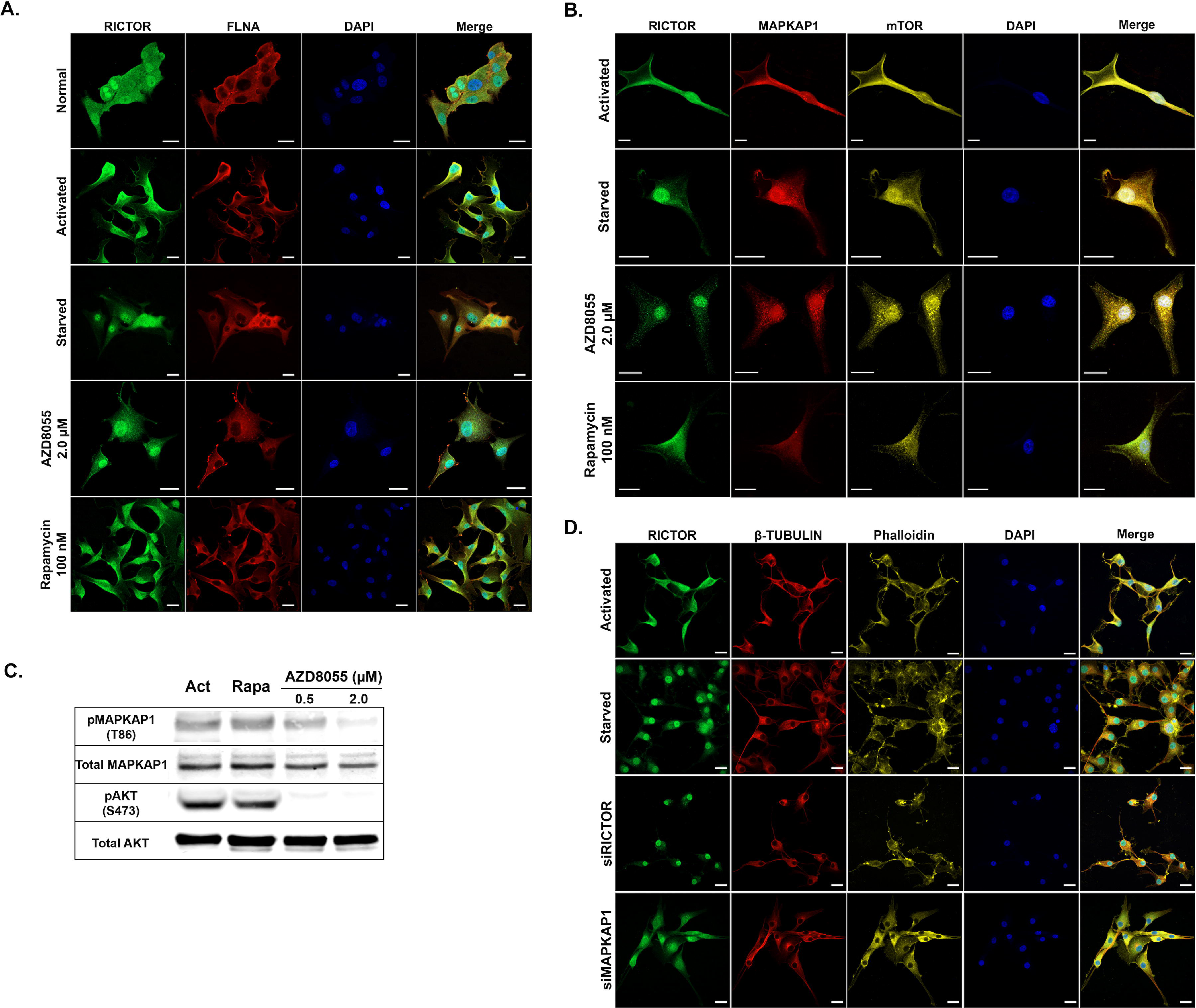
mTORC2 Localization, Integrity, and Effects on Cytoskeleton Rearrangement. (A) Immunofluorescence analysis showing localization of RICTOR and FLNA proteins in U87MG cells under normal, activating, and three inhibition conditions. Scale bar, 20 µm. (B) Immunofluorescence analysis showing localization of RICTOR, MAPKAP1, and mTOR proteins in U87MG cells under activating and three inhibition conditions. Scale bar, 20 µm. (C) Western blot analysis showing mTORC2 complex integrity (pMAPKAP1) and activity (pAKT) in U87MG cells treated with rapamycin and AZD8055 compared to activation condition. (D) Immunofluorescence analysis showing localization of RICTOR and characteristics of F-Actin (phalloidin) and the microtubules (β-TUBULIN) in U87MG cells under activation and knockdown conditions (siRICTOR and siMAPKAP1 treatments). Scale bar, 20 µm.

In addition, the distribution of mTORC2 can contribute to the regulation of its kinase activity. For example, the membrane recruitment of mTORC2 sufficiently promotes AKT phosphorylation (Ebner et al., 2017; Fu and Hall, 2020; Gleason et al., 2019). Additionally, MAPKAP1 was shown to initiate the binding of mTORC2 to the cell membrane (Liu et al., 2015). Thus, we examined the amount of pMAPKAP1 at T86, referring to mTORC2 complex integrity, in U87MG under different treatments (Humphrey et al., 2013; Yang et al., 2015). We detected a similar amount of pMAPKAP1 (T86) in the activated and rapamycin-treated cells but significantly lower in AZD8055-treated cells. Therefore, under migration-promoting conditions, mTORC2 is intact, functionally active, and exists near the cell membrane (Figure 2C). Regarding our hypothesis, we performed IF staining of RICTOR, F-actin, and β-tubulin in U87MG cells. Under an activation condition, an elevated signal of RICTOR is detected near the PM. In contrast, the cells from three inactivation conditions promoted impaired cytoskeleton while RICTOR proteins were absent from the PM area (Figure 2D). These experiments emphasized the importance of active mTORC2 in driving directional motility of GBM cells, leading to the next question through which mechanisms the mTORC2 could directly induce a well-organized cytoskeleton promoting U87MG migration.

### mTORC2 Interactome Supports the Complex’s Function in Brain Cancer Cell Motility

To gain further mechanistic insights into mTORC2-mediated cell migration promotion, we aimed to comprehensively characterize the interacting partners of mTORC2 by the affinity-purification mass spectrometry (AP-MS) method. We decided to use anti-RICTOR antibody coupled with protein A magnetic beads to pull down endogenous mTORC2 and its associated proteins (Figure 3A). To identify specific RICTOR interactors, we performed the quantitative immunoprecipitation combined with knockdown (QUICK) technique (Smits and Vermeulen, 2016) to compare siRICTOR-treated and normal cells.

**Figure 3.**
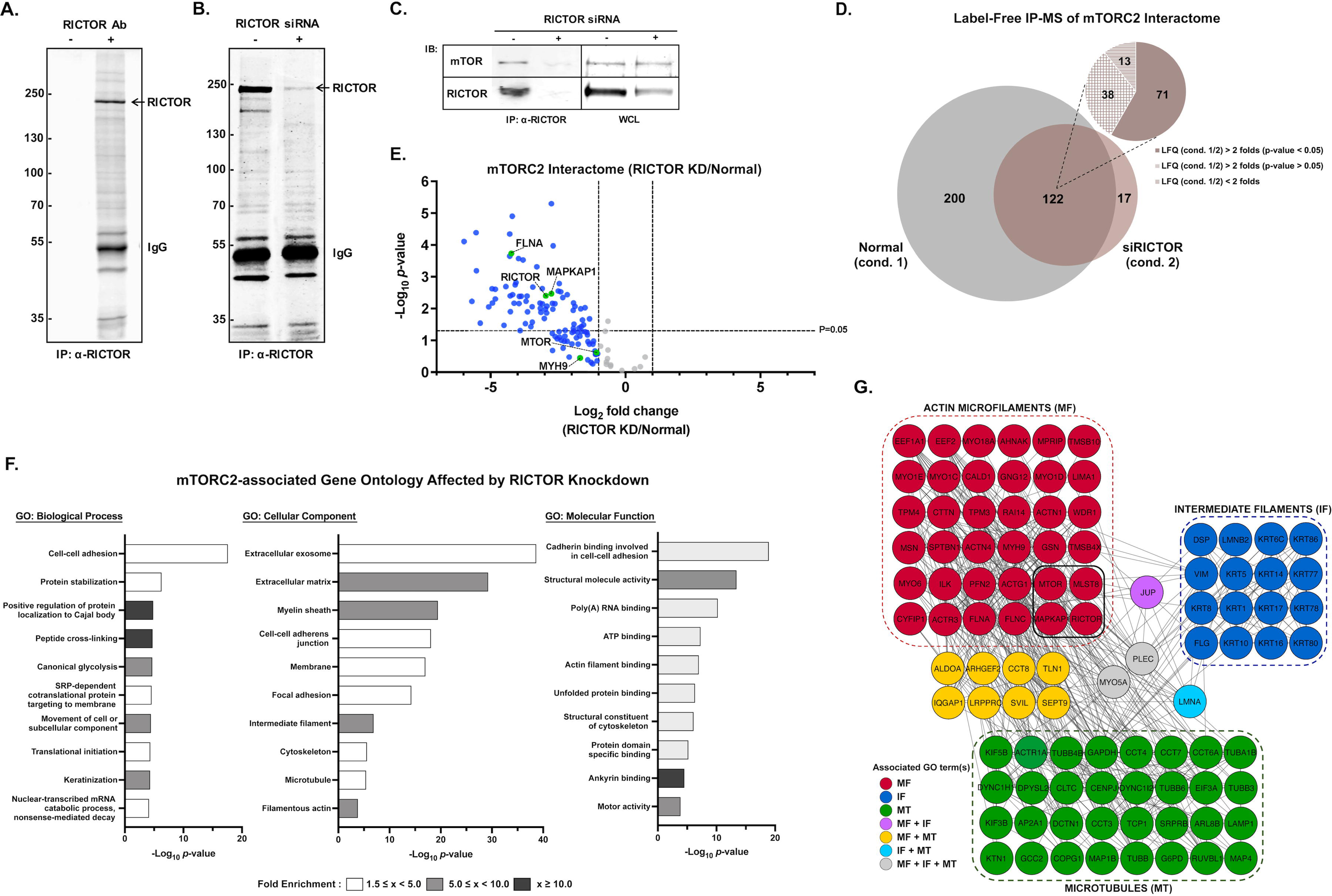
Identification of mTORC2 Interactome by AP-MS. (A) and (B) Coomassie-stained SDS-PAGE gel of an immunoprecipitation assay using protein A magnetic beads incubated with or without anti-RICTOR antibody (A) and RICTOR-IP comparing the control U87MG cells and siRICTOR-treated cells (B). (C) Western blot analysis of immunoprecipitated RICTOR samples and whole-cell lysate (WCL) of U87MG cells treated with RICTOR siRNA compared to control. (D) mTORC2-interacting proteins identified by AP-MS analysis compared between normal and *RICTOR*-knockdown as in (B); n = 3 biological replicates. (E) Volcano plots of results of the AP-MS experiment in (D) showing mTORC2-associated proteins affected by *RICTOR* knockdown. Proteins with fold change (LFQ siRICTOR/normal) ≥ 2 were shown in blue. Known mTORC2 components and interacting proteins were shown in green circles. (F) Gene ontology analysis of 309 proteins negatively affected by *RICTOR* knockdown (decreased by at least 2 folds or absent in siRICTOR-treated IP sample). Ten top-ranked −log_10_(p-value) GOBP, GOCC, and GOMF terms with FDR < 0.05 were shown. (G) mTORC2-interacting protein-protein network based on cytoskeleton-associated proteins from blue circles in (E) affected by *RICTOR* knockdown. Proteins associated with each group of the cytoskeleton were labeled with different colors. Known key components of mTORC2 were in the black-bordered square.

We expected to identify all mTORC2-associated proteins in samples from the normal condition but decreased or absent in the *RICTOR* knockdown (KD) group (Figure 3B). RICTOR and mTOR were reduced in the siRICTOR-treated immunoprecipitated (IP) product, while mTOR amount in the whole-cell lysate was not affected (Figure 3C). We performed label-free quantitative proteomic analysis and reported 309 proteins proximally associated with RICTOR (Table S1). A volcano plot was created from 122 proteins commonly found in both conditions (Figures 3D and 3E). RICTOR amount was depleted by more than seven-folds. We successfully pulled down the known key components of mTORC2, including RICTOR, mTOR, MAPKAP1, and MLST8, as well as FLNA and MYH9, which we reported previously. We also found 17 other known interacting proteins of RICTOR based on the GPS-PROT database (Fahey et al., 2011) as shown in Table S1. We characterized the mTORC2 interactome using DAVID Bioinformatics Resources 6.8 tools. The top-ranked enriched Gene Ontology terms: GOCC, GOBP, and GOMF passing the 5% false discovery rate (FDR) cut-off were selected and shown in figure 3F. Focusing on cell migration regulation, we found that the identified proteins consisted of actin, actin-binding proteins, tubulin isoforms, microtubule-associated proteins, intermediate filaments, and other cytoskeletal regulatory proteins (Figure 3G). Moreover, we could infer that mTORC2 physically interacts with three types of the cytoskeleton and might directly regulate the dynamics of the cytoskeletal networks affecting the migratory phenotypes of GBM cells.

### Quantitative Proteomics Revealed Specific Interacting Partners of RICTOR in U87MG Cells with Various Migratory Ability

Since we observed that mTORC2 altered its intracellular localization when inactivated, we hypothesized that distinct groups of proteins would be highly associated with mTORC2 depending on its activation status. Therefore, we determined all proteins interacting with mTORC2 in two groups of cells with two distinguishable phenotypes, motile and non-motile, using mass spectrometry. The AP-MS experiments were performed using U87MG cells cultured under three different conditions; serum-starvation (ST), serum-activation (ACT), and AZD8055-treated (AZD) conditions. We compared log_2_-transformed label-free quantification (LFQ) intensity of identified proteins from the two pairs: 1) ACT and ST and 2) AZD and ACT. More proteins were identified in ACT eluate than ST and AZD (Figure 4A; Tables S2-S4). We supposed the upregulated and uniquely found proteins in the ACT group would provide more information about how mTORC2 controls cell motility. The LFQ intensity comparisons between the common proteins found in all repeats of each pair of conditions were represented as the volcano plots (Figures 4B and 4C; Tables S5 and S6). Significantly increased proteins of ACT/ST with log_2_ fold change over two include myosin-9 (MYH9), myosin-1D (MYO1D), gelsolin (GSN), and plectin (PLEC), and ATP-binding cassette sub-family F member 2 (ABCF2). Moreover, MYH9, MYO1D, microtubule-associated protein 1B (MAP1B), and actin, cytoplasmic 2 (ACTG1) were significantly decreased in AZD/ACT. Note that GSN and ABCF2 were absent from the RICTOR IP of AZD8055-treated cells (Table S7). We also found that when starved or inhibited by AZD8055, MAPKAP1 was missing from the mTORC2 IP product (Table S7), as previously shown that inactive mTORC2 did not contain MAPKAP1 (Frias et al., 2006; Jacinto et al., 2006; Yang et al., 2006). However, in GBM cells, we observed that mTOR and RICTOR were substantially intact under the inhibitory conditions.

**Figure 4.**
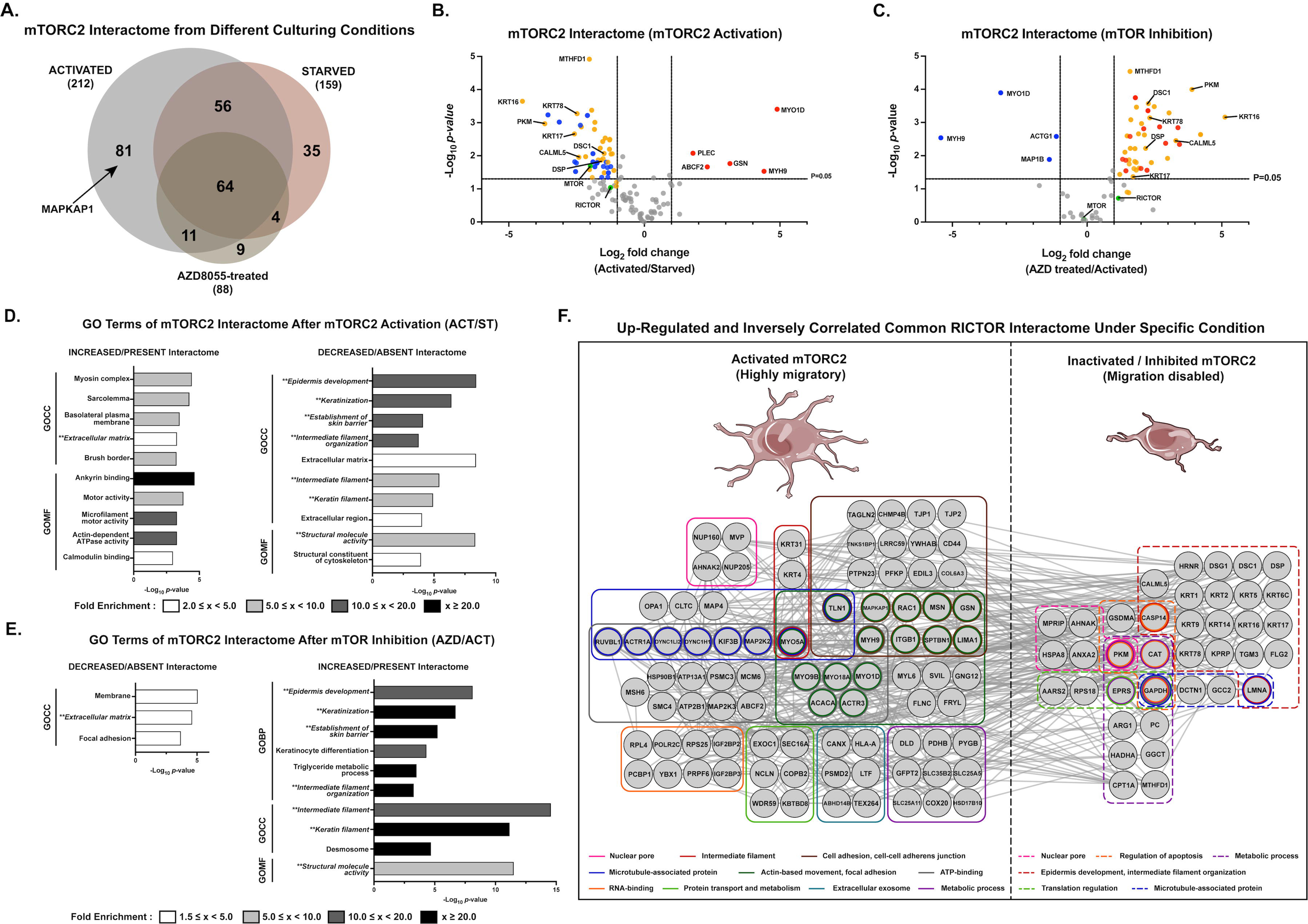
Quantification of mTORC2 Interactome under Cell Migration-associated Conditions. (A) Overlap of mTORC2 interactome identified from AP-MS experiments performed from U87MG cells under activated, starved, and 2.0 µM AZD8055-treated conditions. (B) and (C) Volcano plots of results of the AP-MS experiments in (A) showing changes of mTORC2-associated proteins by serum-stimulated mTORC2 activation in (B) and AZD8055-mediated mTOR inhibition in (C); n=3 biological replicates. Proteins with log_2_ fold change (LFQ ACT/ST or LFQ AZD/ACT) ≥ 1, p < 0.05 were highlighted in red while proteins with log_2_ fold change ≤ −1, p < 0.05 were highlighted in blue. Statistical significance was calculated from LFQ values of each common protein of the two conditions using unpaired two-sided Student’s t-test. Green circles are known as mTORC2-associated proteins. Inversely regulated identical proteins downregulated in ACT/ST and upregulated in AZD/ACT were shown in orange. (D) and (E) Gene ontology analysis of mTORC2 interactome increased/present and decrease/absent by mTORC2 activation (D) and mTOR inhibition (AZD8055-treated) (E). Top-ranked −log_10_(p-value) GOBP, GOCC, and GOMF terms with FDR < 0.05 are shown, and the bar colors represent the fold enrichment of each specific term. (F) Protein-protein interaction network of mTORC2 interactome upregulated in each specific condition with the distinguished phenotype (activated/ highly migratory or inhibited/migration disabled) and found to be downregulated in the opposite condition. All candidate proteins shown in the network were significantly changed with fold change ≥ 2 and p-value < 0.05. Proteins were categorized into various groups labeled by different colors depending on their common associated biological processes, cell components, or functions. Circles with two or more colors represent proteins related to several gene ontologies.

Associated enriched GO terms were annotated to the mTORC2 interactome increased or present after mTORC2 activation (ACT/ST), as well as decreased or absent after the inhibition of mTORC2 (AZD/ACT) (Figures 4D and 4E). We observed that a group of identical proteins (shown as orange circles in volcano plots) was inversely regulated, inferring that these proteins interacted with active mTORC2 and consequently dissociated after mTORC2 inactivation or vice versa (Table S8). Proteins that were statistically less associated with active mTORC2 but tended to bind more to inactive mTORC2 complexes consisted of intermediate filament proteins such as the keratin isoforms and desmosome components. A network of upregulated mTORC2-interacting partners promoting each phenotypic characteristics determining cells’ movement was created (Figure 4F).

### High-grade and low-grade glioma cells have different acquired migration capabilities and RICTOR intracellular localization

Next, we would like to compare the phenotypic changes acquired in the GBM relative to their less invasive counterpart, low-grade glioma. We investigated and compared various characteristics of two cell lines: a non-tumorigenic low-grade glioma cell line (H4) and a glioblastoma (grade IV glioma) cell line (U87MG).

Western blotting analysis showed that U87MG and H4 cell lines expressed comparable amounts of RICTOR. Both cell lines had high mTORC2 activities according to their pAKT (S473) levels, although slightly lower in H4. Effects of AZD8055 and rapamycin on mTORC2 activity were similar (Figure 5A). Then, we performed a cell proliferation assay and observed that H4 cells were as highly proliferative as U87MG cells and were slightly more resistant to AZD8055 (Figure S2A). Hence, the cell proliferation rate of the cell lines could not distinguish brain cancer cells according to their aggressiveness.

**Figure 5.**
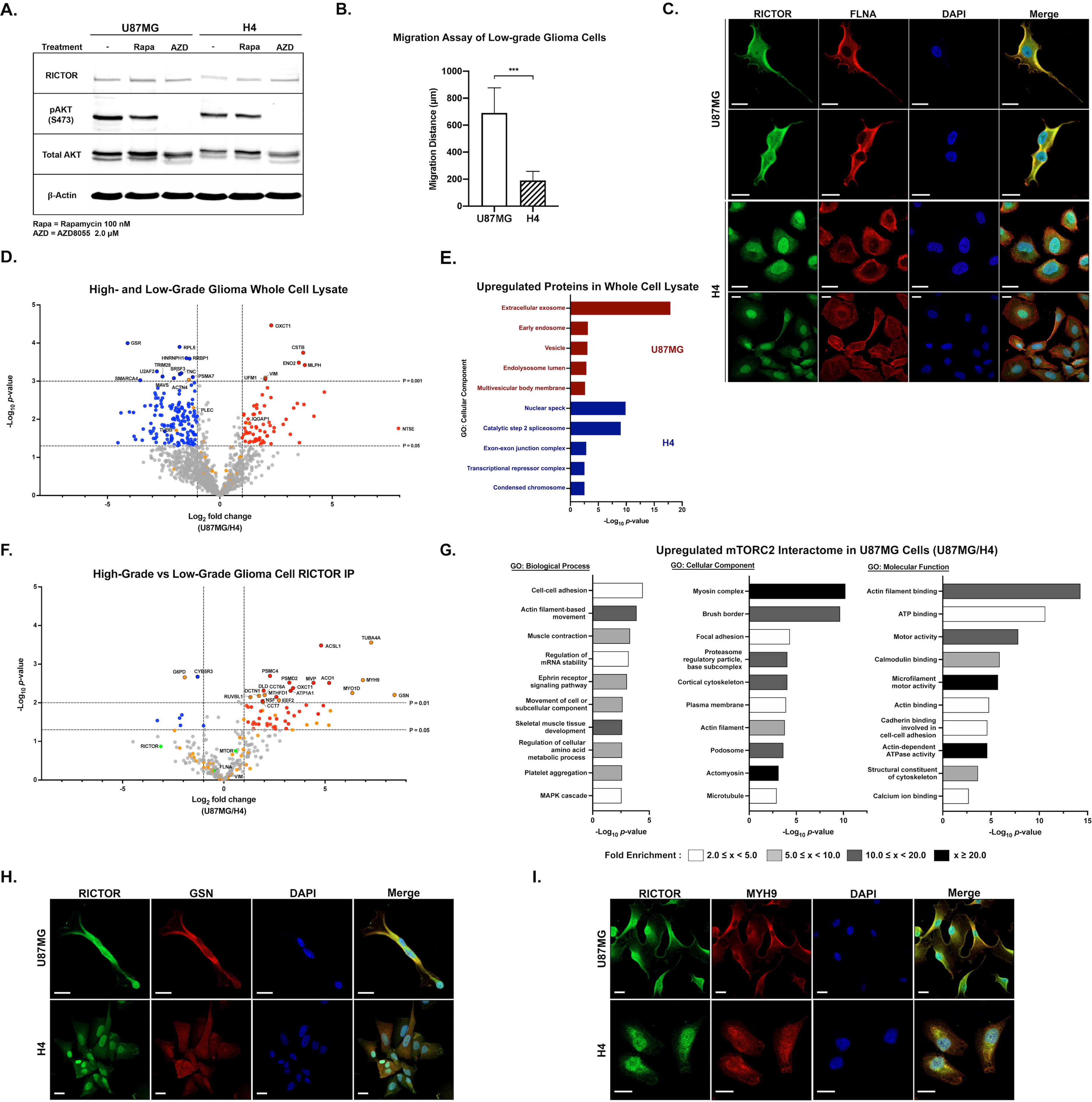
Comparative Analysis of mTORC2 Interactome in High- vs. Low-Grade Glioma Cells. (A) Western blot of RICTOR level and mTORC2 activities (pAKT) using U87MG and H4 cell lysates. (B) Comparison of migration distance between U87MG (experiment shown in figure 1F) and H4 under normal condition; n=3. All data are mean ± SD. ***p < 0.001 (C) Immunofluorescence analysis showing different localizations of RICTOR and FLNA in U87MG and H4 cells. Scale bar = 20 µm. (D) Volcano plots of proteomic analysis from U87MG and H4 whole cell lysates; n=3 biological replicates. Proteins with log_2_ fold change (LFQ U87MG/H4) ≥ 1, p < 0.05 were highlighted in red while proteins with log_2_ fold change ≤ −1, p < 0.05 were highlighted in blue. Proteins with p-value < 0.01 were shown with a black border. Statistical significance was calculated from LFQ values of each common protein of the two conditions using unpaired two-sided Student’s t-test. All cytoskeletal proteins were shown in orange. (E) Gene ontology analysis showing GOCC terms of upregulated and unique proteins of the cell line comparing U87MG and H4. Plots showing ten top-ranked −log_10_(p-value) GOCC terms with FDR < 0.05 of each cell line. (F) Volcano plots of results of the AP-MS experiments in (J) showing changes of mTORC2-associated proteins in U87MG compared to H4 cells; n=3 biological replicates. Proteins with log_2_ fold change (LFQ U87MG/H4) ≥ 1, p < 0.05 were highlighted in red while proteins with log_2_ fold change ≤ −1, p < 0.05 were highlighted in blue. Proteins with p-values < 0.01 were shown with a black border. Statistical significance was calculated from LFQ values of each common protein of the two conditions using unpaired two-sided Student’s t-test. Key mTORC2-interacting proteins were shown in green circles. All cytoskeletal proteins were shown in orange. (G) Gene ontology analysis of increased/present proteins in mTORC2 interactome of U87MG. Plots showing top −log_10_(p-value) GOBP, GOCC, and GOMF terms with FDR < 0.05 and the bar colors representing fold enrichment of each specific term. (H) and (I) Immunofluorescence analysis of RICTOR and GSN (M) or MYH (N) in U87MG and H4 cells. Scale bar = 20 µm.

Moreover, we compared the migration of H4 and U87MG cells (U87MG migration assay was shown in figure 1F). H4 cells migrated approximately three times slower than U87MG (Figure 5B; S2B). We later determined the localization of RICTOR and FLNA inside the H4 cells and found that they were primarily localized around the perinuclear and nuclear region, which is different from U87MG (Figure 5C).

### AP-MS Characterizes Distinct mTORC2 Interactome Profiling of High-Grade and Low-Grade Glioma

Since we expected to pinpoint specific interacting partners prevailing the superior migration of U87MG cells, the label-free quantitative proteomic analyses of H4 and U87MG whole cell lysate were performed. A total of 2733 common proteins were retrieved. For cell line-specific proteins, we identified 523 proteins from H4 and 16 proteins from U87MG. The cytoskeleton-associated proteins exclusively found in each group are listed (Figure S2C). The common 1310 proteins identified from all replicates of both cells were compared and shown in the volcano plot (Figure 5D; Table S9). Significantly upregulated cytoskeletal proteins with over two-fold increase in GBM cells included vimentin (VIM) and IQ motif containing GTPase activating protein 1 (IQGAP1).

Subsequently, we analyzed all upregulated and exclusively found proteins of each group. The top five upregulated GOCC terms of each cell line are shown in figure 5E. We discovered that U87MG expressed more vesicular and extracellular exosome-related proteins, while more nuclear and transcription-related proteins were found in H4. The heatmap compared all common cytoskeletal proteins found in H4 and U87MG cells (Figure S2D). The expression of most cytoskeleton-associated proteins did not show significant differences. We confirmed VIM expression by IF, and the output correlated with our proteomics data that less VIM was observed in H4 than in U87MG (Figure S2E). However, we speculated that the whole cell lysate proteomic results could not specifically answer protein localization and cell migration questions. Therefore, we predicted that observing the mTORC2 interactome would give better insights into the distinct mTORC2-mediated regulatory mechanisms of GBM cell migration.

Consequently, the mTORC2 interactomes of H4 and U87MG were elucidated (Figure S2F; Table S10). The label-free quantification was performed among commonly found proteins, and the log_2_ fold changes of U87MG over H4 were demonstrated (Figure 5F; Table S10). We identified the proteins that elaborated the most remarkable differences between high and low-grade glioma cells. We discovered that gelsolin (GSN), tubulin alpha-4A chain (TUBA4A), myosin-9 (MYH9), and unconventional myosin-Id (MYO1D) had the most considerable fold changes among all mTORC2-interacting proteins (Figure 5F and S2G). Interestingly, GSN, MYH9, and MYO1D had been shown in previous experiments to be highly associated with active mTORC2 in highly migratory U87MG cells and lost their interactions when cells had impaired movement.

Furthermore, we classified all upregulated and specific mTORC2 interactomes in U87MG compared to H4. The associated GO terms consisted of various cytoskeletal structures and activities (Figure 5G). Thus, it could be referred that the interacting proteins more highly associated with mTORC2 in U87MG help promote migrational activities and invasiveness.

Accordingly, we investigated the relationship between GSN or MYH9 and RICTOR in both cell lines. The protein colocalization events between GSN-RICTOR and MYH9-RICTOR were observed in U87MG, but fewer interactions were exhibited in H4. These results were similar to what had been determined in the AP-MS results (Figures 5H and 5I). Overall results suggested that the distinct mTORC2 interactome is one of the key factors supporting the enhanced migration characteristic of GBM cells.

### Gelsolin promotes mTORC2-mediated migration by connecting the complex to actin-binding and membrane-associated proteins to form actin-based structures

Next, we investigated the association between mTORC2 and GSN or MYH9 in GBM cells. The IF stainings of RICTOR, GSN, and MYH9 in the activated, starved, and AZD8055-treated U87MG cells were carried out. We found that RICTOR/GSN/MYH9 colocalized altogether near the PM under the mTORC2-activating condition. In contrast, GSN and MYH9 proteins were observed in the cytoplasm of cells under two mTORC2-inactivating states: starved and AZD8055-treated (Figure 6A and S3A). We later performed *MAPKAP1* knockdown and observed the altered localization of RICTOR to nuclear and cytosolic (Figure 6B). However, the localization of GSN and MYH9 did not appear as immensely altered as RICTOR but tended to be more evenly distributed throughout the cells. Our findings suggested that a functionally active mTORC2 is required for the protein-protein interactions and colocalization between RICTOR/GSN/MYH9 at the PM.

**Figure 6.**
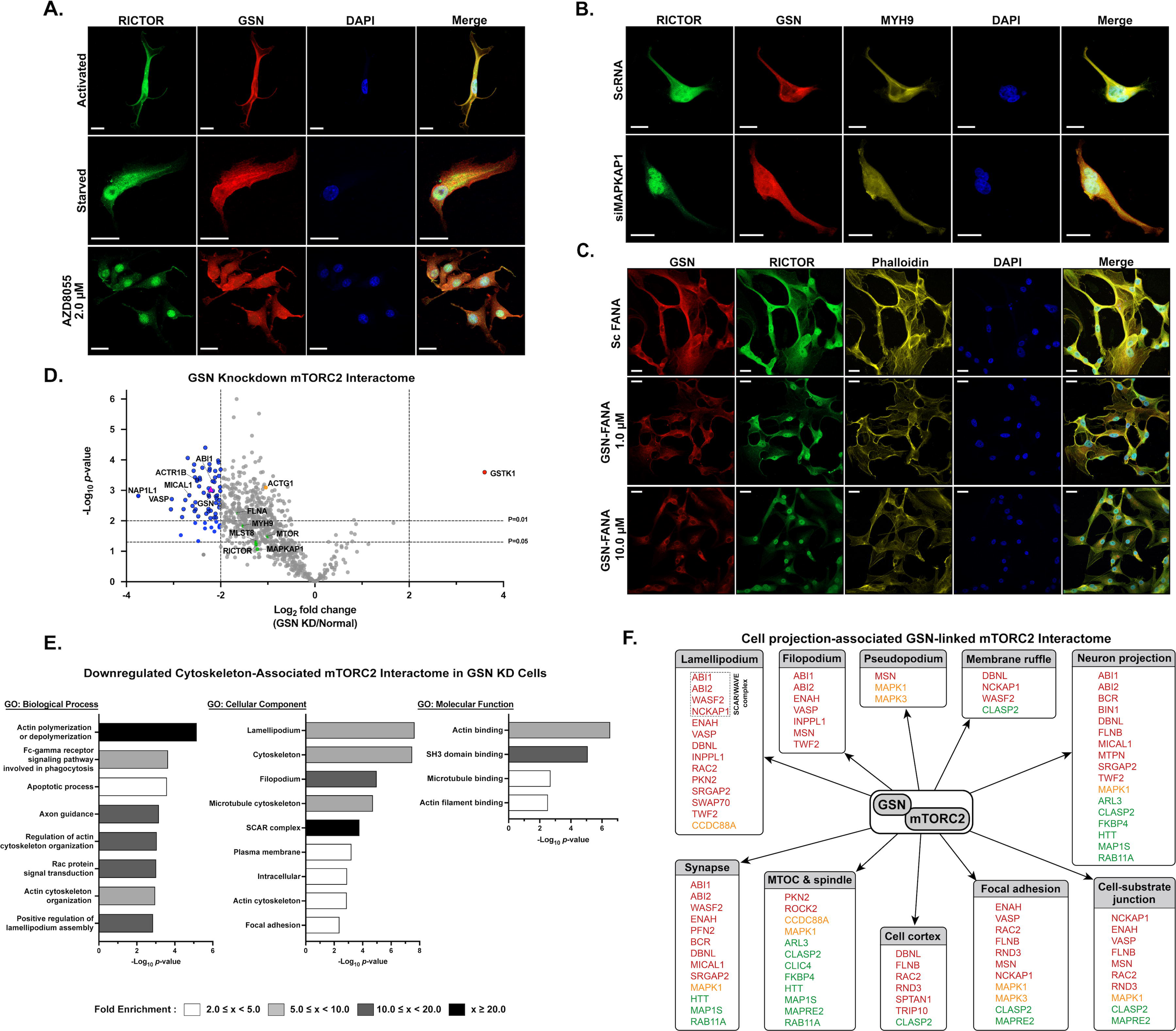
Significance of GSN in mTORC2-Mediated Cell Migrational Control. (A) Immunofluorescence staining of U87MG cells showing localization of RICTOR and GSN (A) under activation, serum-starvation, and inhibition conditions. Scale bar = 20 µM. (B) Immunofluorescence staining of U87MG cells showing colocalization of RICTOR, GSN, and MYH9 of control and *MAPKAP1* knockdown conditions. (Sc: scrambled siRNA). Scale bar = 20 µm. (C) Immunofluorescence staining showing localization of GSN and RICTOR and F-actin arrangement in U87MG cells after *GSN* knockdown (1.0 µM and 10.0 µM of GSN-FANA) compared to scrambled FANA control. Scale bar = 20 µm. (D) Volcano plot of results of the AP-MS experiments in showing changes of mTORC2-interacting proteins; n=3 biological replicates. Proteins with log_2_ fold change (LFQ GSN KD/normal) ≥ 2, p < 0.05 were highlighted in red while proteins with log_2_ fold change ≤ −2, p < 0.05 were highlighted in blue. Proteins with p-value < 0.01 were shown with a black border. Statistical significance was calculated from LFQ values of each common protein of the two conditions using unpaired two-sided Student’s t-test. Key mTORC2-interacting proteins were shown in green circles. GSN was shown in a purple triangle. (E) Gene ontology analysis of decreased/absent proteins in mTORC2 interactome of *GSN* knockdown group. Plots showing top-ranked −log_10_(p-value) GOBP, GOCC, and GOMF terms with FDR < 0.10 and the bar colors representing fold enrichment of each specific term. (F) Diagram showing GSN-linked mTORC2 interactome associated with cell projection structures. (red: MF-associated proteins, green: MT-associated proteins, yellow: MF-MT-associated proteins)

Finally, we comprehensively investigated GSN, the most dynamic protein among all conditions observed. To elucidate the consequences of mTORC2-GSN interaction, we demonstrated the effects of *GSN* knockdown by 2’-deoxy-2’-fluoro-D-arabinonucleic acid antisense oligonucleotides (FANA ASO) against *GSN* gene on the localization of RICTOR, together with the structures of F-actin networks. We found that without GSN, RICTOR distribution was dramatically affected as the proteins were less associated with the PM and only present in the cytoplasm and nuclei. The maximal destructive effect on the F-actin network was determined when cells were treated with 10.0 µM of GSN-FANA (Figure 6C). Overall results suggested that both active mTORC2 complex near plasma membrane and availability of gelsolin in cells could collaboratively promote efficient GBM cell movement.

To characterize the effectors of the mTORC2-GSN connection, we performed a quantitative proteomic analysis of RICTOR IP samples under two conditions: 1) Control and 2) *GSN* knockdown. We aimed to observe all mTORC2-interacting proteins affected by GSN depletion to investigate how mTORC2-GSN interaction can control GBM cell migration. The RICTOR-IP eluates of control and *GSN*-knocked down groups contained partially different proteins (Figure S3C). The common proteins found in both conditions were selected, compared, and shown in the volcano plot (Figure 6D; Table S11).

The results showed that we successfully depleted GSN from mTORC2 pulldown by over four-folds. The knockdown slightly affected the total amount of immunoprecipitated mTORC2 and decreased various mTORC2-associated proteins, especially actin-binding proteins, but did not disrupt the critical components of mTORC2 (mTOR, RICTOR, MAPKAP1, and MLST8). In addition, interactions between mTORC2 and several significant direct interactors such as ACTG1, FLNA, MYH9, MYO1D, TUBA4A, MAP1B, and PLEC were not significantly decreased.

Top-ranked GO biological process terms of the diminished mTORC2-interacting proteins when *GSN* was knocked down were demonstrated (Figure 6E). Furthermore, we focused on the downregulated cytoskeleton-associated mTORC2 interactome and found that the interrupted processes included multiple actin cytoskeleton-related activities. The negatively affected cellular components mainly comprised the cytoskeletal structures supporting cell protrusion. With all the information, we came up with the final network showing effector proteins linked to mTORC2 through GSN or cell projection-associated GSN-linked mTORC2 interactome. (Figure 6F). Our findings help elucidate the PPI network between mTORC2 and specific proteins enabling mTORC2 to regulate the cytoskeletal remodeling and facilitate cell projection.

### Major mTORC2 interactome determines the complex’s functions in invasive brain cancer cells

We proposed the schematic diagram of how active mTORC2 could extensively promote superior cell migration ability of the GBM cells (Figure 7A). When activated, through the linkage with GSN, mTORC2 interacts with many actin-binding proteins and microtubule-associated proteins involved in forming dynamic protrusive structures such as filopodia and lamellipodium. Inversely, inactive mTORC2 complexes interact more with the intermediate filament proteins resulting in a less motile phenotype.

**Figure 7.**
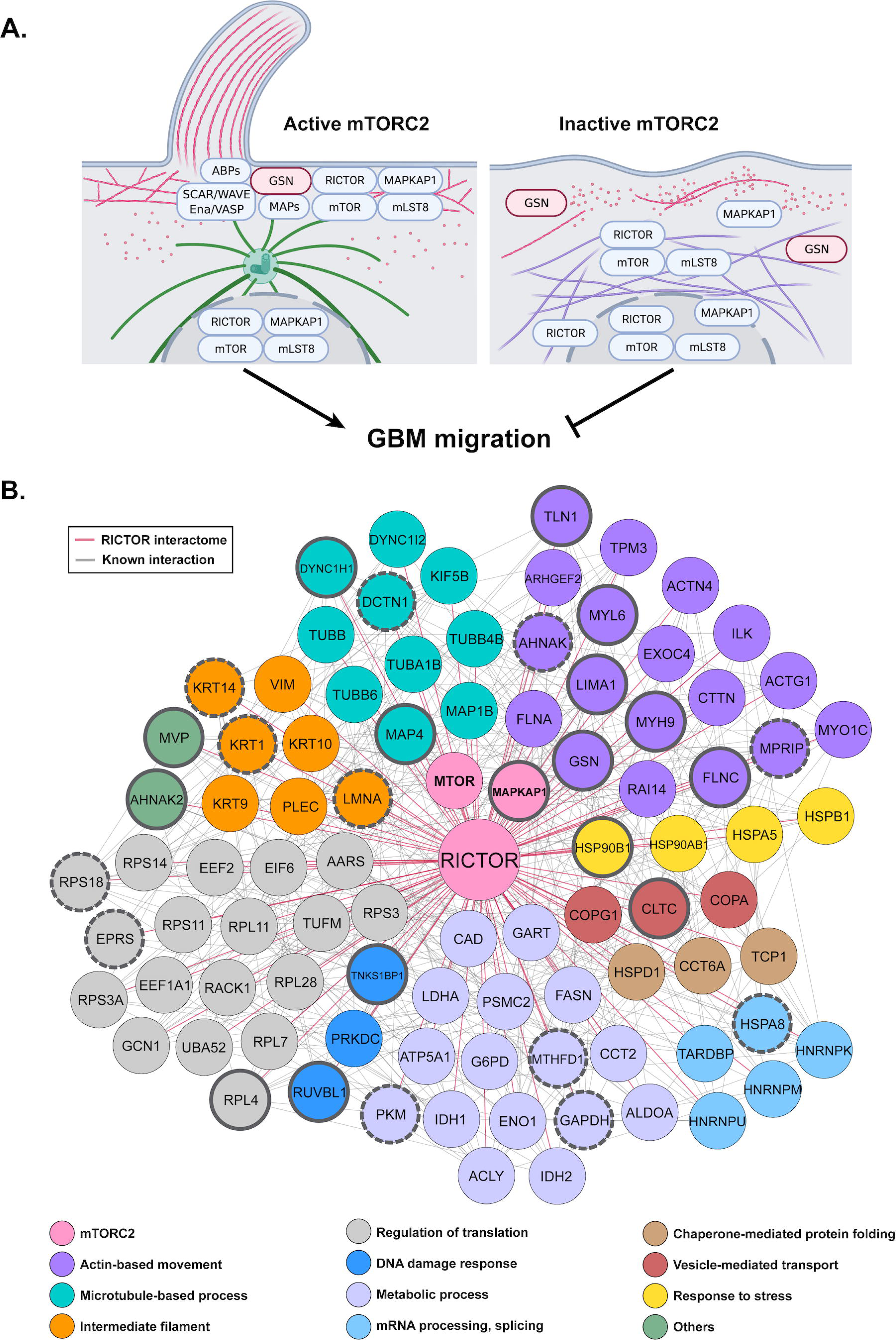
Roles of mTORC2 Interactome in cell migrational control and others. (A) Diagram of the active mTORC2 function in controlling the glioblastoma cell migration via RICTOR-GSN interaction by recruiting actin-binding proteins and microtubule-associated proteins to form cell protrusion structures (left). Under inactivation conditions, mTORC2 loses MAPKAP1 and interacts with the intermediate filament proteins (right). (red: actin cytoskeleton, green: microtubule, purple: intermediate filaments). The graphic was created by Biorender.com. (B) Diagram showing final 86 mTORC2 stable interactome commonly identified from all methods. Proteins were categorized into groups depending on their primary functions and associated biological processes. Pink lines indicate novel interactions. Circles with bold borders represent dynamic mTORC2 interactome upregulated (solid line) or downregulated (dashed line) in migrating glioblastoma cells.

We eventually compared the proteins in the mTORC2 interactome identified by every protein separation and digestion method (Figure S4). There are 101 proteins commonly found in all three groups, including RICTOR, mTOR, and MAPKAP1 (Table S12). All proteins matched with the list obtained from the knockdown experiment were selected and finalized into 88 proteins directly associated with RICTOR (Figure 7B). Apart from the core components of mTORC2, other 86 stable interacting partners of mTORC2 can be categorized into several groups responsible for essential biological processes. To summarize, the results from this study provided more insights into several other aspects of mTORC2 functions in highly invasive glioblastoma, though mainly highlighted the cell migration process. It could further initiate many more investigations on the mechanistic regulation of this multiprotein complex and lead to a better understanding of brain cancer biology.

## DISCUSSION

Glioblastoma multiforme has been depicted as a highly invasive, aggressive, and incurable cancer (DeAngelis, 2001; Neftel et al., 2019; Ohgaki and Kleihues, 2007). This study extensively shows that mTORC2 plays a critical role in GBM migration through the direct regulation of cytoskeleton dynamics. The mTORC2 inhibition by AZD8055 treatment or *RICTOR* knockdown and inactivation by starvation dramatically affects cell migration, actin cytoskeleton, and microtubule organization in GBM cells. Several previous studies have shown that mTORC2 inactivation or ablation could inhibit the migration of many cell types, including cancers (Dada et al., 2008; Daulat et al., 2016; Gulhati et al., 2011; Lamouille et al., 2012; Liu et al., 2010).

Here, we define that RICTOR localization can be used to distinguish glioma cells’ migrational phenotypes in an mTORC2 activity status-dependent manner. Activation of mTORC2 is critically dependent on its key components, RICTOR and MAPKAP1 (Yang *et al*., 2006). The binding of MAPKAP1 initiates the complex integrity of mTORC2 to the PM. Additionally, the association of PH domain of MAPKAP1 with phosphatidylinositol 3,4,5-trisphosphate (PIP3) at the membrane has been shown to regulate mTORC2 (Gan et al., 2011; Liu *et al*., 2015). Ebner et al. have shown that localization of mTORC2 activity can be observed near the plasma membrane, in mitochondria, and some endosomal vesicles (Ebner *et al*., 2017). Also, cytoplasmic and nuclear translocation of mTORC2 components has been determined (Rosner and Hengstschlager, 2008). Even though nuclear or perinuclear mTORC2 function is still elusive (Fu and Hall, 2020; Laribee and Weisman, 2020), our results support the translocation of mTORC2 in GBM occurs when mTORC2 is inactivated, resulting in less plasma membrane-bound and more nuclear and perinuclear mTORC2.

Since the oncogenic signal transduction may differ from the canonical pathway resulting in novel protein-protein interactions (Kovalski et al., 2019), we integrate the quantitative proteomic analysis with modified culturing conditions to vary mTORC2 activation states. This study is the first to describe the dynamic mTORC2 interactome identified from quantitative AP-MS analysis. We demonstrate that mTORC2 activity status defines mTORC2-interacting partners, affecting glioblastoma cells’ migration ability.

Cytoskeletal remodeling is an essential process supporting cancer migration and invasion. Deregulated cellular architecture results in increased formation of various protrusive structures like lamellipodia, filopodia, and invadopodia, affecting cancer cell migration and leading to metastasis (Aseervatham, 2020; Yilmaz and Christofori, 2010). All types of cytoskeleton play pivotal roles in promoting this cancerous behavior (Aseervatham, 2020). Moreover, crosstalk between the actin cytoskeleton and microtubules has been highlighted in cancers (Dogterom and Koenderink, 2019). Our study demonstrates that mTORC2 is one of the master regulators of GBM migration and a central complex connecting numerous cytoskeletal proteins to modulate the cytoskeletal network dynamics by promoting physical interactions between significant players. To elucidate the impact of mTORC2 interactome in enhanced cell migrational control of the highly invasive brain cancer, we have differentially characterized mTORC2 interactome in the GBM compared to non-malignant astrocytoma utilizing the AP-MS technique. The mTORC2-GSN is among the most distinguishable interactions between high- and low-grade cells.

GSN is an important Ca^2+^-dependent regulator of the actin cytoskeleton. It serves as an actin filament severing and capping protein and promotes actin cytoskeleton turnover (Nag et al., 2013). Furthermore, GSN is one of the most abundant actin-binding proteins (ABPs). Its dysregulation has been involved with several pathological conditions in humans, including multiple types of cancers. (Chen et al., 2019; Deng et al., 2015; Kankaya et al., 2015; Li et al., 2012; Liao et al., 2011; Stock et al., 2015). GSN has also been implicated as a regulator promoting cancer cell migration, invasion, and epithelial-mesenchymal transition (EMT) process (Baczynska et al., 2016; Chen et al., 2015; Deng *et al*., 2015; Litwin et al., 2012; Radwanska et al., 2012; Van den Abbeele et al., 2007). Investigation of GSN or gelsolin-like proteins in human gliomas has been reported in several studies as one of the candidate biomarkers for clinical samples (Gdynia et al., 2007; Miyauchi *et al*., 2018; Ohnishi et al., 2009; Yang et al., 2018; Yun et al., 2018). However, the molecular and biological functions of GSN in brain cancer have not been clearly defined.

The molecular mechanism of GSN-mediated actin polymerization and regulation of cell migration is through the binding of phosphatidylinositol 4,5-bisphosphate (PI(4,5)P2), which has gelsolin-uncapping property leading to actin filament elongation (Janmey and Stossel, 1987; Szatmari et al., 2018). GSN severing activity can also be controlled and suppressed by phosphatidylinositol 3,4,5-trisphosphate (PI(3,4,5)P3). Moreover, structural change of mTORC2 located at the PM leading to constitutive activation can be induced by PI(3,4,5)P3 (Fu and Hall, 2020). Loss of PI3Kα, the key PIP3-producing enzyme, increases gelsolin-mediated actin-severing activities (Patel et al., 2018).

Recent studies have described the involvement of the phosphatidylinositol 3,4,5-trisphosphate 5-phosphatase 2 (INPPL1) in cell migration and metastasis of breast cancer and glioblastoma cells. INPPL1 can dephosphorylate PIP3 and PI(4,5)P2 while its inhibition increases glioblastoma migration (Elong Edimo et al., 2016; Ghosh et al., 2018; Ramos et al., 2018). We also identify INPPL1 in the mTORC2 interactome, and its level is affected by the *GSN* knockdown. Our results suggest that mTORC2-GSN interaction might prevent or delay INPPL1 from its catalytic activities. Therefore, one plausible mechanism of how mTORC2 could promote actin polymerization activity is increased PIP2 suppressing GSN-severing activity.

As well as INPPL1, other proteins involved with phosphatidylinositol binding are found to be significantly decreased after *GSN* knockdown, such as coiled-coil domain containing 88A (CCDC88A), profilin-2 (PFN2), twinfilin-2 (TWF2), and phosphatidylinositol-5-phosphate 4-kinase type 2 alpha (PIP4K2A). Interestingly, the CCDC88A (or girdin) has been determined as a critical modulator of the PI3K-mTORC2-AKT signaling pathway that controls cell migration (Lin et al., 2011; Wang et al., 2015; Wu et al., 2016). Hence, the mTORC2 might be primarily involved with the regulation of phosphoinositide signaling. However, more studies are needed to clarify the mechanisms.

Actin polymerization at the PM of the migrating cells promotes the formation of multiple protrusive structures (Litwin *et al*., 2012). We further deconvolute that GSN connects mTORC2 to many actin-binding proteins defined as signaling proteins of actin structures. Our data suggest that when the mTORC2-GSN linkage is disrupted, the most affected processes include actin polymerization/depolymerization, actin-based activities, and actin-microtubule association. The findings conclude that mTORC2 promotes cancer cell migration by inducing the formation of multiple types of actin structures through GSN connection. Both mTORC2 and GSN help promote the PM-localizing process of each other. Additionally, exclusive cytoskeletal proteins in the mTORC2 interactome of U87MG include KIF3B, LIMA1, SVIL, FLNC, and MYO18A. Further studies are essential to clarify how interactions with these proteins lead to advanced GBM migration.

Finally, we identify 86 proteins that are mTORC2’s stable interactors in GBM cells and highlight the significance of GSN. With GSN, mTORC2 can be constantly localized close to the plasma membrane. The complex would also perform its known functions by activating the PKCs and FLNA to generate highly dynamic networks. The stable mTORC2 interactome identified in our study could provide additional information regarding other biological roles of mTORC2 in brain cancer.

In summary, this study provides the novel fundamental idea of a specific regulatory mechanism of aggressive brain cancer cell migration through the relationship between mTORC2 and GSN. Additional players and other mechanistic regulations promoting glioblastoma migration ability await further investigations, which could be deciphered using quantitative proteomics and systems biology approaches.

## ACKNOWLEDGEMENTS

This research is supported by the Ratchadaphiseksomphot Endowment Fund (RGN-2559-059-14-30), Part of the “Research Grant for New Scholar CU Researcher’s Project” Chulalongkorn University; Ratchadaphiseksomphot Endowment Fund (CU59-005-HR) of Chulalongkorn University; the Research Fund for DPST Graduate with First Placement (Grant no. 026/2015), The Institute for the Promotion of Teaching Science and Technology (IPST), Thailand; the Research Grant for New Scholar (Grant no. MRG6180215), Thailand Research Fund (TRF) and the Office of the Higher Education Commission (OHEC); Research Grant for New Scholar (Grant no. RGNS 63-007), Office of the Permanent Secretary, Ministry of Higher Education, Science, Research and Innovation.

## AUTHOR CONTRIBUTIONS

N.C., P.W., and T.P. participated in the study design. N.C. and N.P. performed cell-based and imaging experiments. N.C., P.W., and N.K. performed mass spectrometry experiments. N.C. and P.W. carried out the quantitative proteomic and bioinformatic analyses. N.C. and P.W. wrote the manuscript. P.K., C.A., T.P., F.T., and J.A.L. provided suggestions.

## CONTACT FOR RESOURCE SHARING

Further information and requests for resources and reagents should be directed to and will be fulfilled by the Lead Contact, Naphat Chantaravisoot.

## EXPERIMENTAL MODEL

### Cell Lines

U87MG and H4 cells were maintained in low glucose Dulbecco’s modified Eagle’s medium (DMEM) supplemented with 10% (vol/vol) fetal bovine serum (FBS), 1% (vol/vol) penicillin/streptomycin at 37°C with 5% (vol/vol) CO_2_. Both cell lines were purchased from ATCC. For an activation condition, cells were starved in the serum-free media for 24 hours before adding the DMEM with 10% FBS back for another 24 hours. The starvation condition was performed by replacing the regular media with serum-free media for 24 hours before the experiment.

## METHOD DETAILS

### Drug Treatment

U87MG and H4 cells were cultured in low glucose DMEM containing 10% (vol/vol) FBS, 1% (vol/vol) penicillin/streptomycin at 37°C with 5% (vol/vol) CO_2_ until they reached 80% confluency. The cultured cells were serum-starved in serum-free DMEM for 24 hours before a 24-hour treatment of different concentrations of rapamycin or AZD8055 in regular media. For live-cell imaging, U87MG cells in drug-treated groups were serum-starved for 24 hours before starting the experiment. Then, drug-containing and serum-supplemented DMEM was added to the cells for another 24 hours during the imaging process.

### Gene Knockdown Experiment

*RICTOR* knockdown experiments were performed using On-TARGETplus Smartpool Human RICTOR siRNA or On-TARGETplus Non-targeting Control Pool (Dharmacon) were transfected into U87MG cells using Lipofectamine^TM^ 3000 Transfection Reagent (ThermoFisher) in Opti-MEM media (Gibco). *MAPKAP1* knockdown experiments were performed using Accell Human MAPKAP1 siRNA or Accell Non-targeting Control Pool (Dharmacon). *GSN* knockdown experiments were performed using 2’-deoxy-2’-fluoro-D-arabinonucleic acid antisense oligonucleotides (FANA Antisense Oligos; FANA ASOs) against *GSN* or scrambled control FANA purchased from AUM BioTech.

U87MG cells were treated with each RNAi and their scramble controls for 24 or 48 hours before the experiments were performed, depending on assay types. The cell lysate was collected 48 hours after siRNA treatment for the following experiments, including western blotting, immunofluorescence staining, and immunoprecipitation. For live-cell imaging and wound-healing assay, U87MG cells were transfected with RICTOR siRNA for 24 hours before the start and incubated for another 24 hours while performing the experiments.

### Western Blotting Analysis

U87MG and H4 cells were seeded to 6-well plates until reaching 80% confluency before the different treatments were performed. For protein extraction, cultured cells were lysed with a lysis buffer containing 1% Triton X-100, 150 mM NaCl, 20 mM Tris HCl (pH 7.4), 1mM EDTA, and EDTA-free protease inhibitor cocktail (PIC) (Roche). Cells were lysed on ice for 15 minutes and centrifuged at 16,000 x g for 10 minutes. Total protein concentrations in the whole cell lysate supernatants were determined by Bradford protein assay (Bio-Rad). The protein extracts from various samples (25-35 ug) were equally loaded into the SDS-PAGE gel, separated by electrophoresis, and transferred to a nitrocellulose membrane (Bio-Rad). The membranes were blocked in Odyssey® Blocking Buffer (TBS) (LI-COR) for at least 1 hour at room temperature or overnight at 4°C, then probed with primary antibodies overnight at 4°C. Membranes were washed in TBST and then incubated with IRDye® secondary antibodies (LI-COR). The membranes were scanned on the Odyssey® CLx Imaging Systems (LI-COR). The following antibodies were used: anti-phospho-S6 (S235/236), anti-S6, anti-phospho-AKT (S473), anti-AKT, anti-phospho-MAPKAP1(T86), anti-mTOR, anti-phospho-FLNA (S2152) (Cell Signaling Technologies), anti-FLNA (EMD Millipore), anti-RICTOR, anti-MAPKAP1 (Abcam).

### Cell Proliferation Assay

The cell proliferation of U87MG and H4 was observed using CellTiter 96^®^ AQ_ueous_ One Solution Proliferation Assay (Promega). Cells were cultured in a 96-well plate for 72 hours under activation and AZD8055 treatment conditions. The absorbance at 490 nm was measured to determine the cell viability of each well. The average intensity to the control well ratios at 24, 48, and 72 hours after the drug treatment were plotted onto the graphs using GraphPad Prism.

### Immunofluorescence

U87MG and H4 cells (1×10^4^ cells per well) were cultured in 8-well chamber slides (Lab-Tek). Before staining, cells were starved, activated, or treated with AZD8055 for 24 hours. For the knockdown conditions, cells were incubated with siRICTOR, siMAPKAP1, FANA-GSN, and scramble controls for 48 hours prior to the steps. Cells were fixed with 4% Paraformaldehyde, lysed with 0.2% Triton X buffer, blocked with 1% BSA, and incubated with primary antibodies (anti-RICTOR, anti-MAPKAP1, anti-GSN, anti-β-tubulin (Abcam), anti-mTOR, anti-VIM (Cell Signaling Technologies), anti-FLNA (Millipore), Alexa Fluor 647 Anti-MYH9 (Abcam), Alexa Fluor 488-phalloidin (Invitrogen)) overnight at 4°C. Samples were stained with secondary antibodies, including Alexa Fluor 488, Alexa Fluor 594, Alexa Fluor 647 Anti-Rabbit antibodies, and Alexa Fluor 568 Anti-mouse antibody. Zenon Rabbit IgG labeling kit (Invitrogen) was used to be directly conjugated to primary antibodies when necessary. Nuclei of cells were stained with DAPI. Slides were mounted with ProLong anti-fade mountant (Invitrogen). Cells were visualized by an LSM800 with an Airyscan confocal microscope (Zeiss).

### Live-Cell Imaging of Cytoskeleton

To visualize the actin cytoskeleton and the microtubules, U87MG cells were labeled by silicon rhodamine jasplakinolide targeting actins (SiR-Actin) and silicon rhodamine Docetaxel targeting microtubules (SiR-Tubulin) (Spirochrome) at 200 nM for 3 hours prior to the imaging processes. Time-lapse microscopy started at hour 0 before the treatments were administered or at hour 24 of the siRNA-treated group for 16-24 hours. Images of fluorescently labeled U87MG were taken to observe the cytoskeletal network using time-lapse mode at 15-minute intervals. Differential Interference Contrast (DIC) images were overlayed and combined into merged pictures. All pictures were taken by an LSM800 with an Airyscan confocal microscope (Zeiss).

### Cell Migration Assay

#### Wound-Healing Assay

U87MG cells were grown in 12-well plates in a single layer until they became over 90% confluent. One scratch per well was made using a small plastic pipette tip, creating an approximately 1000 µm-wide gap. Images of each sample group were taken at hour 0 before drug administration. Cells treated with RICTOR siRNA were pre-incubated with the siRNA for 24 hours before the first image. Cells were activated, starved, and treated with AZD8055 2.0 µM or rapamycin 100 nM, and images were acquired again after 6, 12, and 24 hours.

#### Live-Cell Imaging

Live-cell imaging of cell migration assay was performed using DIC mode of LSM800 Airyscan confocal microscope (Zeiss). Images had been continuously taken from hour 0 (right after drug administration) for 14-24 hours using time-lapse imaging. Cell migration tracking was performed using TrackMate tool on Fiji program (Tinevez et al., 2017). Single-particle tracking mode was used to follow U87MG cell migration coordinately. Tracking lines were generated, and each colored line represents the migration path of a single glioblastoma cell.

The migration recovery experiment compared the 24-hr starved cells and three conditions of 24-hr siRICTOR treated U87MG cells. The cell migration was observed with time-lapse microscopy for 18 hours. The reduced serum media containing RICTOR siRNA in two siRICTOR-treated groups were half or entirely replaced with regular media (low glucose DMEM with 10% FBS). Live-cell imaging was performed to investigate the recovery of migration ability of RICTOR-depleted cells after being supplemented with the serum to activate mTORC2.

### Cell Lysate Proteomics

U87MG and H4 cells were cultured in 150-mm tissue culture dishes. Cells were harvested and lysed with 8M urea, 100 mM Triethylammonium bicarbonate (TEAB), 1X EDTA-free PIC (Roche), and 1X phosphatase inhibitor (Thermo Scientific™) and homogenized for 5 minutes. Lysates are centrifuged at 16,000 × g for 5 minutes at 4°C. Protein concentrations were determined by BCA assay (Thermo Scientific™). Protein concentration was adjusted to 10 mg/mL using lysis buffer. Proteins were reduced with 10 mM dithiothreitol for 30 minutes at 37°C, subsequently alkylated in the dark with 40 mM iodoacetamide for 45 minutes at 25°C, and quenched with 10 mM dithiothreitol for 15 minutes at 25°C. Samples were diluted with 100 mM TEAB until containing 0.6 M urea and digested with sequencing grade modified trypsin (Promega) using a 1:150 enzyme-to-substrate ratio at 37°C for 16 hours. The digested samples were acidified with 100% trifluoroacetic acid (TFA), with the final sample solution containing 0.5% TFA. Then, tryptic peptides were cleaned up with reversed-phase C18 solid-phase extraction (SPE) columns. The peptide concentrations were further determined by Quantitative Fluorometric Peptide Assay (Thermo Scientific™ Pierce™). Finally, the peptides were dried using SpeedVac (Thermo Scientific™)

### Affinity Purification Coupled with Mass Spectrometry

#### Immunoprecipitation of mTORC2

U87MG and H4 cells were cultured in 150-mm dishes under normal, mTORC2-activating and mTORC2-inactivating conditions mentioned previously for 24 hours before being harvested. Cells were collected after washing with PBS containing 1X EDTA-free PIC (Roche) and stored at −80°C for subsequent experiments. Endogenous proteins were studied to avoid artifact effects from cells overexpressing epitope-tagged proteins that might further ambiguously affect the PPIs and to understand all physical interactions under various culturing conditions with minimal confounding factors. To affinity purify mTORC2, we followed our original protocol with partial modifications (Chantaravisoot *et al*., 2015). Cells were lysed in lysis buffer (2% CHAPS, 50 mM HEPES, pH 7.4, 150 mM NaCl, 1mM Na_3_VO_4_, 1X EDTA-free PIC (Roche)). Cleared supernatant after centrifugation (16,000 × g for 10 min) was mixed with anti-RICTOR antibody (Abcam) overnight at 4 °C. After that, SureBeads™ Protein A magnetic beads (Bio-rad) were mixed with the lysate for 60 minutes at room temperature for affinity purification. The beads were collected and washed five times with wash buffer containing high salt (0.1% CHAPS, 50 mM HEPES pH 7.4, 300 mM NaCl, 2 mM DTT) to remove non-specifically bound proteins. The proteins were eluted from the beads by mixing with Laemmli buffer and boiled at 95°C for 10 min. Protein samples were run in 8-10% SDS-PAGE. Gels were stained using Imperial Protein Stain solution (Thermo Scientific™) to visualize protein bands. Western blotting was also performed to confirm the key components of purified mTORC2.

To investigate the true mTORC2 interactome in U87MG cells, we adopted the idea of QUICK (quantitative immunoprecipitation combined with knockdown) technique (Selbach and Mann, 2006) and slightly modified it by performing label-free quantitative proteomic analysis instead of SILAC labeling. Experiments were performed in five replicates comparing siRICTOR-treated cells to normal U87MG cells. Co-immunoprecipitated proteins were separated by SDS-PAGE followed by in-gel digestion to identify the IP products.

#### Mass Spectrometry Sample Preparation

##### In-Gel Digestion

The immunoprecipitated mTORC2 samples were fractionated by SDS-PAGE using 8% Tris-Glycine gel. The gel was stained with Imperial Protein Stain solution (Thermo Scientific™). After that, the whole gel was washed with distilled water on a glass plate, and each lane was excised into eight slices for MS analysis. Each gel slice was further cut into small pieces (1 mm^3^), then washed in 50% acetonitrile (ACN) in 25 mM triethylammonium bicarbonate (TEAB) buffer overnight to be fully destained and washed once again with the same buffer. All gels were dried down by vacuum centrifugation. Proteins in the gel slices were reduced with 10 mM dithiothreitol (DTT) at 56°C for 1 hour, followed by alkylation with 50 mM iodoacetamide at room temperature for 45 min. Tryptic digestion in 25 mM TEAB was performed using sequencing grade trypsin (Promega) overnight at 37 °C. Peptides were eluted with 50% ACN, and 1% trifluoroacetic acid (TFA), dried with a SpeedVac and stored at −20 °C before LC-MS/MS analysis.

#### GELFREE Fractionation and In-Solution Digestion

The mTORC2 interactome from U87MG cells under three different conditions (starvation, activation, and inhibition) was immunoprecipitated as previously described, followed by the fractionation step using the gel-eluted liquid fraction entrapment electrophoresis (GELFREE) method (Tran and Doucette, 2008) to ensure that all samples were equally fractionated for label-free quantitative proteomic analyses. Then, the samples were digested using the eFASP in-solution digestion protocol with trypsin (Erde et al., 2014). For GELFREE separation, IP eluates were loaded into an 8% cartridge of the GELFREE 8100 system (Expedeon). Proteins were subsequently separated into 6 fractions. The collected samples were processed by eFASP protocol using a 30 kDa MWCO cut-off centrifugal filter (Millipore). Laemmli buffer was exchanged into 0.2% deoxycholic acid (DCA) in 20 mM ammonium bicarbonate (ABC). Proteins were trypsin-digested on the filter after reduction and alkylation. DCA was removed by acid precipitation using TFA and three rounds of ethyl acetate extraction. Peptides were dried with vacuum centrifugation and stored at −20 °C.

#### On-Bead Digestion

To compare the mTORC2 interactome affected by gelsolin knockdown, the pulled-down proteins were digested by on-bead digestion technique modified from previous literature (Keilhauer et al., 2015). We aimed to identify as many associated interacting partners as possible, including multiple layers of interactions. After the purification step, additional wash steps were performed with the beads using wash buffer I (0.05% IGEPAL CA-630, 150 mM NaCl, 50 mM Tris HCl pH 7.5, and 5% glycerol), followed by wash buffer II (150 mM NaCl, 50 mM Tris, and 5% glycerol) to remove trace amount of detergents. During the elution step, 25 μL of elution buffer I (2 M Urea, 50 mM Tris, and 1 mM DTT, 5 ng/µL trypsin) was incubated with the beads for 30 minutes for the partial digestion to release mTORC2 interactome from the antibodies. Proteins were then eluted and alkylated twice with 50 μL elution buffer II (2 M Urea, 50 mM Tris, and 5 mM Chloroacetamide) for 30 minutes. Finally, the supernatants of each sample were pooled in a new tube and incubated overnight at room temperature for complete digestion. The digestion was stopped by adding 2 μL trifluoroacetic acid (TFA). The final peptide clean-up step was performed using C18 StageTips (Empore) with Jupiter 300 (Phenomenex). Peptides were dried by SpeedVac (Thermo Scientific™) and resuspended in 0.1% formic acid for LC-MS/MS analysis.

#### Mass Spectrometry LC-MS/MS

Tryptic peptides were analyzed by Thermo EASY-nanoLC™ 1000 System equipped with a 25-cm PepMap C18 EASY-Spray column and Q-Exactive Plus Orbitrap mass spectrometer (Thermo Scientific™) with a flow rate of 300 nL/min over a 90-minute gradient. The sample injection volume was 10 µL. Solvent A was 0.1% formic acid (FA) in water, and solvent B was 0.1% FA in acetonitrile. The gradient was ramped from 5 to15% solvent B for 45 minutes, then increased from 15 to 40% within 8 minutes, then jumped to 95% in 2 minutes, and remained at 95% for 5 minutes. Eluted peptides from nano-LC were sprayed into the orbitrap mass spectrometer at 2.5 kV capillary voltage in the positive ion mode and 255 °C capillary temperature. MS/MS was operated at Top 10 data-dependent acquisition (DDA). For the full MS scans, 75,000 resolution, target ion at 1 x 10^6^, and maximum IT of 120 ms were chosen. For the MS/MS mode, 12,500 resolution, 3 x 10^4^ AGC, and 25 ms maximum IT were set. An isolation window was set to 1.2 m/z. The normalized collision energy was used at 27. The dynamic exclusion was set at 30 seconds with dynamic fixed first mass.

#### LC-MS/MS Data Processing

The proteome database search was performed by MaxQuant 1.6.2.1 with Andromeda algorithm (Cox and Mann, 2008; Cox et al., 2011) against human SwissProt proteome and common contaminants (http://www.thegpm.org/crap/) databases. The search allowed up to 2 missed cleavages with a mass tolerance of 10 ppm and 0.02 Da for MS1 and MS2, respectively. Carbamidomethylation (C) as static modification, oxidation (15.994915@M), phosphorylation (79.966331@S, T, Y) and ubiquitination (114.042927@K) as variable modifications, were included. The second peptide, dependent peptide, and match between runs were enabled. The match between runs feature was set at 0.7 min time window, 20 min alignment window. The false discovery rate (FDR) was controlled at 1%.

#### Label-Free Quantitative Proteomic Analysis

Label-free quantitation (LFQ) by spectral counting was enabled without normalization in MaxQuant 1.6.2.1 program in the experiments requiring quantitative analyses. Statistical analysis was performed by Perseus 1.6.0.2 software (Tyanova et al., 2016) with no imputation. All LFQ intensities involved in the bioinformatics analysis are log_2_-transformed. Proteins identified by site and reverse matches were removed. To quantify the relative changes of each identified protein, the spectral counting value of each condition was compared to another condition. First, we expected to identify all mTORC2-associated proteins present in normal IP samples but appeared to be over two-fold decreased or absent in the *RICTOR* knockdown IP samples of at least three out of five replicates.

To compare mTORC2 interactome between the two cellular conditions: starvation and activation, or activation and AZD8055-treated. We also performed label-free quantitative proteomic analysis. Only proteins detected in all three repeats passed our filtering criteria for selected candidates. Proteins that were significantly increased or decreased by at least two folds (*p*-value < 0.05) or unique (absent in two or all repeats of the comparing group) under certain conditions might be correlated with specific mTORC2 functions responsible for cells’ migratory phenotypes.

In the proteomic analysis comparing U87MG and H4 proteins, log_2_ LFQ values of the candidate proteins found in three biological replicates were used to generate heatmaps using Java TreeView Cluster 3.0 with average linkage clustering. All proteins on the list acquired significant p-values (p<0.05) when compared between the two groups. The cell lysate heatmap was generated from cytoskeletal proteins commonly found in the lysates. In addition, the IP heatmaps showed common mTORC2-associated cytoskeletal proteins identified from the immunoprecipitated samples of U87MG and H4 cells.

Lastly, to identify GSN-linked mTORC2-associated proteins, we immunoprecipitated mTORC2 from GSN-knockdown cells compared to wild-type U87MG cells. In this case, we prepared the AP-MS samples using the on-beads digestion method to reduce the loss of GSN-associated mTORC2-interacting proteins co-purified with RICTOR. To consider which proteins are linked to mTORC2 through GSN, we set up the filtering criteria that the amount of each candidate protein should be decreased in the GSN-knocked down condition at least relatively equal to or less than GSN itself (log_2_ fold change GSN KD/normal < −2) when compared to the control group. We supposed these specific proteins could not physically link to mTORC2 without GSN.

### Protein-Protein Interaction Network Construction

#### Gene Ontology Analysis

We characterized the mTORC2 interactome using DAVID Bioinformatics Resources 6.8 tools (Huang da et al., 2009a; b), then annotated the mTORC2-associated gene ontologies by comparing our protein list to glioma proteome as a background in the enrichment analysis to avoid tissue-specific enriched terms. The background list includes 6359 proteins derived from a combination of all identified proteins from every mass spectrometry experiment that we performed using glioma cell lines (U87MG and H4), including both whole cell lysate and immunoprecipitated samples. To compare mTORC2 IP under different treatment conditions, we used the background list of 2083 proteins identified from all immunoprecipitated experiments under any treatment conditions to narrow it down to only highly significant GO terms related to the activation or inhibition of the mTORC2 pathway. Every protein in the background list must be found in at least three replicates. After performing the analysis, we reported the most significant candidate GO terms. All selected GO terms must have high confidence passing 5% or 10% FDR. The protein network interactomes were constructed in Cytoscape v3.8.0 software (Shannon et al., 2003). The known protein-protein interactions among all candidate RICTOR interacting partners were imported using GeneMANIA App based on protein colocalization and physical interactions. Redundant edges were removed, and novel edges were highlighted.

## QUANTIFICATION AND STATISTICAL ANALYSIS

The data represented in the graphs are mean ± SD. The number of experiments (n) shown in the figure legends refers to biological replicates or experimental replicates, depending on the appropriateness. All statistical analyses were performed using GraphPad Prism 9. Significance of all statistical analyses was considered significant when passing the 95% confidence level (* p < 0.05, ** p < 0.01, *** p < 0.001, **** p < 0.0001). The unpaired two-tailed Student’s t-test was used when comparing two independent groups. A two-way ANOVA and Ordinary one-way ANOVA with Tukey’s multiple comparisons test were used to compare multiple groups with single or multiple parameters. For gene ontology, false discovery rate (FDR), p-values, and fold enrichment values were calculated by DAVID Bioinformatics Resources 6.8 tools (Huang da *et al*., 2009a; b).

## DATA AND SOFTWARE AVAILABILITY

The accession number for the mass spectrometry data reported in this paper is PXD035256. Raw data have been deposited to Mendeley Data and are available at https://doi.org/10.17632/4dmy59w389.1.

